# Cochlin-Expressing Memory B (COMB) Cells Are Enriched in Autoimmune Diseases and Display a Distinct Activation Profile

**DOI:** 10.1101/2025.03.24.644109

**Authors:** Becca Belmonte, Mohini Gray

## Abstract

Rheumatoid arthritis (RA) is the most common chronic autoimmune arthritis, causing joint damage and affecting multiple organs over time. B cells play a key role in driving the disease by producing autoantibodies, releasing cytokines, and presenting antigens to T cells. While B cell depletion therapies can help reduce inflammation, they remove all CD20+B cells indiscriminately, which can increase infection risk and interfere with important regulatory immune functions.

To better define pathogenic B cell subsets, we performed single-cell RNA and ATAC sequencing on circulating and synovial B cells from patients with early, untreated RA. We identified a novel population of memory B cells expressing cochlin (COCH), termed cochlin - expressing memory B (COMB) cells. While detectable at low levels in healthy controls, COMB cells are significantly expanded in RA and exhibit a distinct transcriptional profile indicative of immune activation. Epigenetic analysis showed that COMB cells have more open chromatin at key immune regulatory regions, including sites bound by NF-κB family transcription factors like RELA and REL. This suggests that these cells are primed to respond to inflammatory signals. We also looked at publicly available datasets and found COMB-like cells in the blood of people with systemic lupus erythematosus (SLE) and Sjögren’s syndrome (SjS), as well as in inflamed kidney tissue from patients with SLE. COMB cells across these diseases share a conserved gene expression signature, pointing to a common memory B cell programme associated with autoimmunity.

These findings define COMB cells as a previously unrecognised, transcriptionally and epigenetically distinct memory B cell subset enriched across autoimmune diseases, offering new insights into B cell–mediated pathology and potential therapeutic targets.

## Introduction

Autoimmune diseases are rising globally, with the annual incidence and prevalence increasing by 19.1% and 12.5%, respectively [1]. These conditions may soon dominate as leading medical disorders, causing severe organ damage and life-threatening complications [2]. The economic burden is immense, with annual treatment costs exceeding $70 billion USD and climbing, placing a significant strain on already stretched healthcare systems.

Current treatments often fail to achieve sustained remission, carry serious side effects, and inadequately target underlying disease pathways. Indeed, despite advances in biologic therapies for autoimmune diseases such as rheumatoid arthritis (RA), they offer few advantages over optimised combinations of synthetic DMARDs [3]. Moreover, therapies for conditions such as Sjögren’s syndrome (SjS) and systemic sclerosis are largely ineffective at altering disease outcomes. This underscores the urgent need for deeper insights into autoimmune pathogenesis to develop more effective, safer and targeted therapies.

B cells are central to autoimmunity, driving disease through autoantibody production, antigen presentation, cytokine secretion, and by controlling ectopic lymphoid structure formation [4]. Specific subsets of B cells, such as double-negative (DN) 2 B cells, characterised by a CD11c^+ve^T-bet^+ve^CD27^-ve^IgD^-ve^ phenotype, are expanded in systemic lupus erythematosus (SLE) [5] as well as RA, where they serve as precursors to antibody-secreting cells (ASCs) in the inflamed rheumatoid synovium [6]. These cells can bypass germinal centres via extrafollicular responses to produce short-lived ASCs, forming immune complexes that activate innate immune cells, driving chronic inflammation [7].

Commonly people with one autoimmune disease will also manifest another. For example, up to a third of patients with RA also have SjS, whilst autoimmune thyroid disease is twice as common in RA patients compared to the general population [8]. In addition, SLE often coexists with SjS [9, 10]. What characterises all these conditions is B cell dysfunction leading to autoantibody production, which suggests there may be overlapping mechanisms driving them [11]. Defects in B cell tolerance, hyperactive signalling pathways, and cytokine dysregulation (e.g., elevated BAFF) may all contribute to the persistence and overactivity of autoreactive B cells. These mechanisms may be triggered by multiple mechanisms including genetic susceptibility, environmental irritants (such as smoking), or other stochastic factors. They are then amplified by autoimmune T cells, which are themselves sustained by the enhanced antigen-presenting function of autoimmune B cells, creating a positive feedback loop of inflammation.

B cell-depleting therapies (BCDT), that target CD20 expressed on the B cell surface have shown efficacy in RA, SLE, and multiple sclerosis (MS) [4]. However, non-specific depletion of all CD20+ B cells has been linked to increased morbidity and mortality in SARS-CoV-2-infected patients [12]. In addition, gold-standard synthetic disease modifying drugs such as methotrexate, are poorly tolerated by many patients. This highlights the dual role of B cells, as both drivers of chronic autoimmune inflammation and as key components of immune defence. Furthermore, B cell depletion can potentially worsen autoimmunity by removing regulatory B cells, disrupting self-tolerance to apoptotic cells, and reducing the production of anti-inflammatory cytokines, including IL-10 [13-15]. These limitations underscore the need for more selective strategies that target pathogenic B cells while preserving protective and regulatory immune functions.

Single-cell sequencing and advanced analysis techniques have transformed our ability to study human autoimmunity. These tools allow us to examine individual cells within complex populations, providing detailed insights into cellular states, gene expression, and functional diversity. This approach helps identify rare cell populations, trace cellular lineages, and discover new therapeutic targets. Combining clinical expertise with hypothesis-driven research and sophisticated data analysis, offers the best hope of uncovering the complex cellular and molecular mechanisms driving human autoimmune diseases.

Using this approach, we analysed B cell populations across multiple autoimmune diseases, looking specifically for shared populations of B cells that were enhanced in autoimmunity. In the circulating B cells of newly diagnosed, untreated RA patients, we identified a distinct subset of pathogenic B cells, which we termed **<u>CO</u>**chlin expressing **M**emory **B** (COMB) cells. A key characteristic of COMB cells was their expression of cochlin, a protein previously believed to be primarily located in the cochlea of the inner ear. This subset of B cells was also enriched in the circulation of patients with SLE, and SjS. Notably, COMB cells were enriched in the rheumatoid synovium, where they represented the dominant memory B cell population. They were also present in renal lupus tissue. These findings suggest that COMB cells may function as a major subset of B cells in autoimmune rheumatic diseases.

## Materials and Methods

### Patients

The use of human samples was approved by the South East Scotland Bioresource NHS Ethical Review Board (Ref. 15/ES/0094) and the use of healthy donor blood was approved by AMREC (Ref. 21-EMREC-041). Informed consent was obtained from all study participants prior to sample collection. The patient characteristics are described in Table 1.

**Table 1:** summarises the distribution of B cells across the eight samples along with clinical data.

| Sample Name | Number of B Cells sequenced | Patient age | Sex | Disease Activity Score in 28 joints (DAS 28) | ACPA titre (AU/ml) |
| --- | --- | --- | --- | --- | --- |
| OA 1 | 1396 | 69 | Female | Not applicable (N/A) |  |
| OA 2 | 532 | 72 | Female | N/A |  |
| OA 3 | 3056 | 57 | Female | N/A |  |
| OA 4 | 838 | 63 | Female | N/A |  |
| RA 1 | 820 | 64 | Female | 7.58 | >340 |
| RA 2 | 1775 | 71 | Female | 7.1 | 158 |
| RA 3 | 1732 | 42 | Female | 5.5 | >340 |
| RA 4 | 2800 | 71 | Female | 5.43 | 129 |

### Cell isolation

Peripheral blood mononuclear cells (PBMCs) were isolated from total blood using Histopaque-1077 separation according to the manufacturer’s protocol (Sigma-Aldrich). The PBMCs were collected, washed in PBS, and counted. PBMCs were resuspended in resuspension media (RPMI-1640 (Gibco) supplemented with 2 mM l-glutamine, 100 U mL-1 penicillin, and 100 μg mL-1 streptomycin, with 40% fetal calf serum (FCS)) then an equal volume of freezing media (resuspension media supplemented with 30% dimethyl sulfoxide (DMSO, Fisher Reagents)) was added to give final density of 10×10^6^ cells mL^−1^ in 15% DMSO. The sample was then frozen to −80°C in a Mr Frosty Freezing Container (ThermoFisher Scientific) then stored in liquid nitrogen until further use.

### BD Rhapsody run and sequencing

PBMC samples were thawed and resuspended. Briefly, cryovials were removed from liquid nitrogen and thawed rapidly in a water bath at 37°C. The cells were sequentially diluted 5 times with warm media with 1:1 incremental volume addition added in a dropwise manner, gently swirling between each volume. The cell suspension was centrifuged at 230xg for 6 minutes then washed in PBS. After defrosting, cells were labelled with sample tags and AbSeq Ab-Oligos according to the manufacturer’s protocol (BD Biosciences). Cell suspensions were then stained in FACS buffer (PBS supplemented with 1% FCS) at room temperature for 20 minutes with fluorescently conjugated antibodies as described in Supplementary Table 1. Cells were washed and re-suspended in FACS buffer. Single CD19+ve DAPI-ve CD3-ve B cells were then sorted with the FACS Fusion Q and FACS Diva 8.0.1 (BD Biosciences). The full gating strategy is shown in Supplementary Fig. S1. Immediately after sorting, one OA and one RA sample were mixed in equal ratios. The cell suspension was loaded onto the BD Rhapsody cartridge and libraries were prepared according to the manufacturer’s protocol (BD Biosciences). The number of cells per sample loaded onto the BD Rhapsody cartridge are shown in Supplementary Table 1. Whole transcriptome and AbSeq libraries were sequenced using the NextSeq 2000 at the Genetics Core at the Edinburgh Clinical Research Facility.

### Data pre-processing

The FASTQ files were processed using the BD Rhapsody Sequence Analysis Pipeline (BD Rhapsody) and aligned to the GRCh38 reference genome using the SevenBridges BD Rhapsody Pipeline. The R data file was loaded into R (v4.1.0) using the Seurat [16] (v4.4.1) package.

### Clustering, dimensionality reduction, and differential expression

Unsupervised clustering based on the first 20 principal components of the most variably expressed genes was performed using Seurat::FindNeighbours and Seurat::FindClusters with the resolution 0.8. Clusters were visualized using the uniform manifold approximation and projection (UMAP) method and annotated by analysing the expression of canonical cell markers and SingleR [17]. Differential gene expression between clusters was examined using the Wilcox test using Seurat::FindMarkers and Seurat::FindAllMarkers. Results of these tests were presented using Seurat::DotPlot.

### Trajectory inference

Trajectory inference was performed using destiny [18] (v3.18.0) and monocle [19] (v1.3.7). Destiny uses diffusion maps while monocle uses reverse graph embedding to infer pseudotime.

### Cell abundance

For both scRNA-seq and scATAC-seq, differential abundance of cell clusters was calculated using miloR [20] (v2.2.0). Milo calculates neighbourhoods using a nearest neighbour graph and tested using a negative binomial generalized linear model.

### Module scores and gene set enrichment analysis

Module scores were created using the Seurat::AddModuleScore function. The top DEGs for COMB cells with >2 log2FC and an adjusted p-value <0.01 were then used as the input list for Seurat::AddModuleScore creating a module score for each cell. The cells were then visualized using the UMAP method and contour plots layered over to show the cells within the top 5% for each module score.

Gene lists that are specific for synovial, blood RA, and blood OA COMB cells were found using Seurat::FindMarkers and the top genes with a log2FC > 2 and an adjusted p-value < 0.01 were selected. Genes that were in all three gene lists were used as the input list for Seurat::AddModuleScore creating a module score for each cell.

Raw data from human PBMCs from Primary Sjogren’s Syndrome patients was downloaded from GSE214974 [21]. Raw data from human PBMCs from SLE patients was downloaded from GSE142016 [22] and GSE135779 [23]. Data from SLE renal tissue was downloaded from the ARK portal. The data was input and integrated into individual objects by disease and tissue type. B cells were selected using SingleR [17] annotation. Cells with the number of features between 100 and 3000, and with <10 percent mitochondrial reads were kept for further analysis.

The results published here are in whole or in part based on data obtained from the ARK Portal (https://arkportal.synapse.org/). This work was supported by the Accelerating Medicines Partnership® Rheumatoid Arthritis and Systemic Lupus Erythematosus (AMP® RA/SLE) Program. AMP® is a public-private partnership (AbbVie Inc., Arthritis Foundation, Bristol-Myers Squibb Company, Foundation for the National Institutes of Health, GlaxoSmithKline, Janssen Research and Development, LLC, Lupus Foundation of America, Lupus Research Alliance, Merck & Co., Inc., National Institute of Allergy and Infectious Diseases, National Institute of Arthritis and Musculoskeletal and Skin Diseases, Pfizer Inc., Rheumatology Research Foundation, Sanofi and Takeda Pharmaceuticals International, Inc.) created to develop new ways of identifying and validating promising biological targets for diagnostics and drug development Funding was provided through grants from the National Institutes of Health (UH2-AR067676, UH2-AR067677, UH2-AR067679, UH2-AR067681, UH2-AR067685, UH2-AR067688, UH2-AR067689, UH2-AR067690, UH2-AR067691, UH2-AR067694, and UM2-AR067678).

Gene set enrichment analysis (GSEA) was carried out using clusterProfiler::enrichGO [24] (v3.0.4). Gene Ontology (GO) gene sets were downloaded from org.Hs.eg.db (v7.5.1). The differentially expressed genes identified by the venn diagram were ranked by log2 fold change and used as the input. The GSEA results were visualized with clusterProfiler::dotplot and clusterProfiler::cnetplot.

### 10X scATAC-seq run and sequencing

PBMC samples were thawed and resuspended. Briefly, cryovials were removed from liquid nitrogen and thawed rapidly in a water bath at 37°C. The cells were sequentially diluted 5 times with warm media with 1:1 incremental volume addition added in a dropwise manner, gently swirling between each volume. The cell suspension was centrifuged at 230xg for 6 minutes then washed in PBS. Cell suspensions were then stained in FACS buffer (PBS supplemented with 1% FCS) at room temperature for 20 minutes with fluorescently conjugated antibodies as described in Supplementary Table 1. Cells were washed and re-suspended in FACS buffer. Single CD19+ve DAPI-ve CD3-ve B cells were then sorted with the FACS Fusion Q and FACS Diva 8.0.1 (BD Biosciences). The full gating strategy is shown in Supplementary Fig. S1. Immediately after sorting, nuclei were isolated from sorted cells following the manufacturer’s protocol (10X Genomics). scATAC-seq libraries were prepared using the Chromium Single Cell Platform and the ATAC Library Kit V2 (10X Genomics). Libraries were then sequenced using the NextSeq 2000 at the Genetics Core at the Edinburgh Clinical Research Facility.

### scATAC-seq data pre-processing

The FASTQ files were processed using Cell Ranger ATAC pipeline (10X Genomics) and aligned to the GRCh38 reference genome. The raw peak matrices were loaded into R (v4.4.1) using the Seurat [16] and Signac [25] packages.

Each sample was pre-processed separately before integration. Cells that had fewer than 3,000 peaks, more than 30,000 peaks, had fewer than 15% of reads within peaks, had a nucleosome signal below 4 or had a TSS enrichment greater than 3 were removed to exclude low-quality cells or technical artifacts. The gene activity matrix was added to the object using Signac::GeneActivity. Contaminating non-B cells were then excluded by selecting cells annotated as B cells with SingleR [17].

### scATAC-seq clustering, dimensionality reduction, and differential expression

Following quality control, the cells were clustered using the functions Signac::RunTFIDF, Signac::FindTopFeatures, Signac::RunSVD, Signac::FindNeighbors, and Signac::FindClusters, with the LSI components 2-50 at a resolution of 0.6. The scRNA-seq data was then projected onto the scATAC-seq data using Signac::FindTransferAnchors and Signac::TransferData. Clusters were annotated according to the resulting predicted labels, along with canonical gene expression. Clusters were visualized using the uniform manifold approximation and projection (UMAP) method. Differential gene expression between clusters was examined using the logistic regression test using Seurat::FindMarkers and Seurat::FindAllMarkers. Results were visualized using ggplot2 [26].

### Motif analysis

A list of motif frequency matrices was downloaded from the JASPAR database (v0.99.10) and added to the Seurat object using Signac::AddMotifs. Motifs that were specific for COMB cells were identified using differentially accessible regions (DARs) that were enriched in COMB cells. The top regions were selected from DARs with a log2FC > 2, an adjusted p-value < 0.05, and were expressed in over 20% of COMB cells. These regions were used as input for the Signac::FindMotifs function and results were visualized with Signac::MotifPlot. Motif activity score was also calculated per cell using chromVar [27] (v1.26.0). Activity of selected motifs were visualized as features using Seurat::FeaturePlot.

Direct targets of c-Rel as listed in [28] were used as inputs to calculate module scores using Seurat::AddModuleScore. Results were visualized using Seurat::FeaturePlot and ggplot2 [26] (v3.5.1). Differences in module scores between clusters was calculated using a linear mixed effects model with the R package “lme4” [29], with the original sample as a random effect. Multiple comparisons were performed with Tukey using the R package “multcomp” [30].

## Results

### Developmental Trajectory and Population Analysis of Circulating B Cell Subsets in RA and OA Patients

To determine whether patients with RA exhibit unique patterns of circulating B cell expression, we first studied patients with newly diagnosed, active seropositive RA who had not yet started disease-modifying therapy. We compared them to patients with the degenerative, non-autoimmune condition called osteoarthritis (OA).

To gain an unbiased understanding of all circulating B cell subsets, we performed single -cell RNA sequencing on 12,949 B cells from RA and matched OA patients. Using the R package Seurat, we identified ten distinct clusters: five naïve, four memory, and one antibody-secreting cell (ASC) cluster (Fig. 1A). Among the naïve B cell subsets, five were distinguished based on differential gene expression, collectively accounting for just over half of the total B cells (Table 2).

**Table 2:** Distribution of B Cell Subsets in RA and OA patients by Single-Cell RNA Sequencing

| Cluster | Number of Cells total | % Cells total |
| --- | --- | --- |
| Pre-B | 124 | 1.0 |
| Early B | 1395 | 10.8 |
| Transitional | 1477 | 11.4 |
| Naïve | 4853 | 37.5 |
| Early Activation | 1285 | 9.9 |
| Non-switched Memory | 1113 | 8.6 |
| Switched Memory | 768 | 5.9 |
| COMB | 1265 | 9.8 |
| DN2 | 547 | 4.2 |
| ASC | 122 | 0.9 |

**Fig 1.**
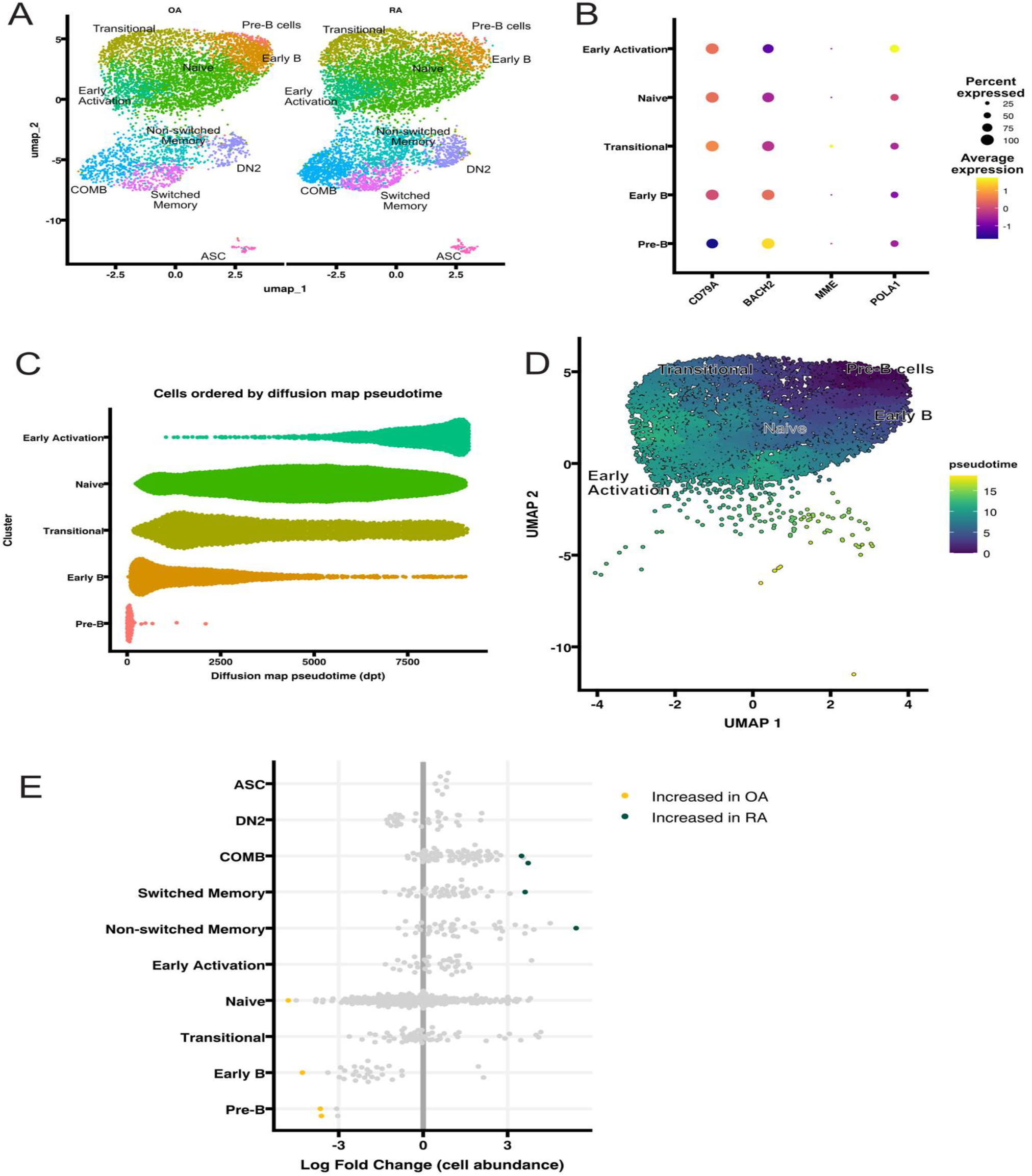
Developmental Trajectory and Population Analysis of B Cell Subsets in RA and OA Patients. **1A**. UMAP visualization comparing B cell populations between OA (left) and RA (right) samples. Distinct populations are identified, including Pre-B cells, Early B cells, Transitional B cells, Naive B cells, Early Activation B cells, Non-switched Memory cells, Switched Memory cells, DN2 cells, COMB cells, and antibody-secreting cells (ASCs). **1B**. Dot plot illustrating differential gene expression across B cell subsets. Dot size represents the percentage of cells expressing each gene, while colour indicates average expression levels. Key markers include *CD79A, BACH2, MME*, and *POLA1*. **1C**. Diffusion map pseudotime analysis showing the developmental progression of B cell populations. Naïve subsets are listed on the y-axis and pseudotime is shown on the y-axis. Each cell is represented as a dot. The trajectory progresses from Pre-B cells (red) through Early B (orange), Transitional (yellow), Naive (green), to Early Activated (turquoise) cells. **1D**. UMAP visualization with pseudotime overlay demonstrating the developmental progression of B cell subsets. The colour gradient from purple (early pseudotime) to green (late pseudotime) indicates progression over time. Cell populations are labelled to show their developmental relationships. **1E**. Differential abundance analysis comparing cell populations between RA and OA samples. Log fold changes in cell abundance are shown for each subset. Neighbourhoods with a p-value < 0.05 are shown in colour, with yellow indicating significant increases in OA and green indicating increases in RA.

We sequenced 12,949 B cells and clustered them using Seurat into 10 clusters, with 5 naïve clusters, 4 memory clusters, and ASCs (Fig 1A). To our knowledge, there has not yet been a study focused on unbiased peripheral B cell subset identification, so we performed a detailed examination of all B cell subsets present in our data. The naïve populations were identified using differential expression analysis to find marker genes for each population. All naïve B cell clusters were classified as *IGHD*+ and *CD27*-. Pre-B cells were identified by low *CD79A* expression, a component of the cell surface BCR complex [31, 32], indicating that these cells have decreased BCR expression (Fig 1B). Pre-B cells also express high levels of *BACH2*, which is correlated with the initiation of immunoglobulin (Ig) gene rearrangements and is associated with negative selection at the pre-B cell receptor checkpoint [33, 34] (Fig 1B). Early B cells had decreased expression relative to pre-B cells, but still high expression of *BACH2*, while also having relatively increased expression of *CD79A*, indicating that they have passed the pre-B cell checkpoint and are upregulating surface BCR expression (Fig 1B). Transitional B cells expressed *MME* (CD10) most highly, consistent with canonical transitional B cell markers [35, 36] (Fig 1B). The subset labelled naïve B cells were *IGHD* and *IGHM* positive, and *CD27* and *MME* negative, consistent with a canonical naïve B cell phenotype [37] (Fig 1B). The early activation population highly expressed *POLA1*, the catalytic subunit of DNA polymerase *α*, suggesting that these cells are undergoing increased replication and potentially proliferation (Fig 1B).

To further define the naïve populations and their relations to each other, we calculated the diffusion map of naïve cells and used that to infer pseudotime, using the R package “destiny”. This shows that pre-B cells are the earliest population, followed by early B cells, transitional, naïve B cells, and finally early activation cells (Fig 1C). Trajectory analysis using monocle supported this, with pre-B cells at the earliest time point, and early activation cells being the latest in pseudotime (Fig 1D). These results agreed with the cluster identification results using differential expression.

We then investigated how abundant all populations were in OA and RA patients using miloR. This analysis creates cell neighbourhoods within our dataset and calculates if each neighbourhood has equal proportions of cells from each disease, or if the neighbourhood is skewed towards one condition. We found neighbourhoods within the pre-B, early B, and naïve cell clusters that were significantly enriched in OA patients (P<0.05) (Fig 1E). Conversely, we identified neighbourhoods within memory B cell clusters that were significantly enriched in RA patients (P<0.05) (Fig 1E). This suggests that circulating B cells are more likely to be activated and antigen-experienced in RA when compared to OA. We then focused on the memory and ASC populations.

### Characterization of Memory B Cell Populations, Developmental Trajectories, and COMB Cells in RA and OA

We characterized memory B cells and antibody-secreting cells (ASCs) based on differential gene expression. All memory and ASC populations were CD27-positive (Fig 2A). Non-switched memory cells expressed higher levels of *IGHD* and *IGHM* compared to other memory subsets, with lower levels of *IGHG1* and *IGHG4*, indicating they had not undergone class switching. Most memory populations were CD27-high, except for DN2 cells, which were CD27-low and exhibited high expression of *LILRB2* and *ITGAX*, consistent with known DN2 markers [5, 6] (Fig 2A). Switched memory and COMB cells were *IGHD*/*IGHM*-low, reflecting class switching. COMB cells were distinguished from other switched memory subsets by increased expression of *IGHE* and *COCH*. Due to their high *COCH* expression, we named these cells <u>C</u>ochlin expressing <u>m</u>emory B (COMB) cells. ASCs were identified by their high expression of immunoglobulin genes (*IGHG1, IGHG4*) and CD27 (Fig 2A).

**Fig 2.**
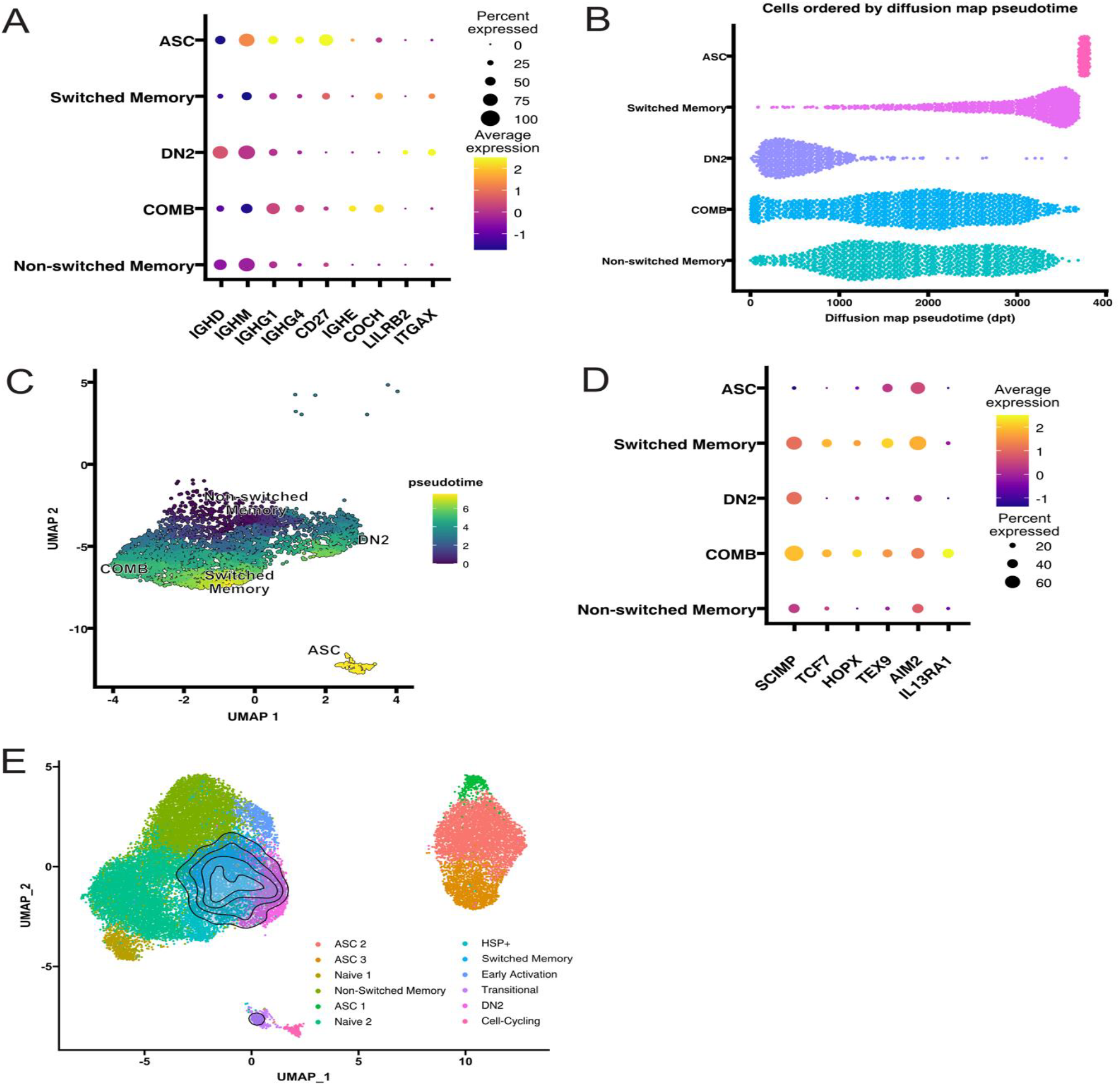
Characterization of Memory B Cell Populations and Their Developmental Trajectories in RA and OA. **2A**. Dot plot showing differential gene expression across memory B cell populations, including non-switched memory, switched memory, DN2, COMB, and ASCs. Dot size represents the percentage of cells expressing each gene, while colour indicates average expression levels. Key genes include *IGHM, IGHG1, IGHG4, CD27, COCH, LILRB2*, and *ITGAX*. **2B**. Diffusion map pseudotime analysis illustrating developmental progression of memory B cell populations. Memory subsets are listed on the y-axis and pseudotime is shown on the y-axis. Each cell is represented as a dot. The trajectory begins with non-switched memory cells and progresses through switched memory subsets, DN2 cells, COMB cells, and culminates in ASCs. **2C**. Monocle UMAP visualization coloured by pseudotime progression, showing spatial organization of memory B cell populations. The gradient represents developmental progression from non-switched memory subsets through switched memory subsets and DN2 cells to ASCs. **2D**. Dot plot illustrating expression of key memory-associated genes (*SCIMP, TCF7, HOPX, TEX9, AIM2*, and *IL13RA1*) across memory B cell populations. Dot size represents the percentage of expressing cells, while colour indicates expression levels. **2E**. UMAP projection of scRNA-seq data from 27,053 synovial B cells from RA patients (n=3) with the module scores for COMB cells, the contour overlay represents the position of the top 5% of module scores. These contour lines overlap significantly with the population previously identified as switched memory B cells in the RA synovium.

To investigate developmental relationships, we performed diffusion map pseudotime analysis on memory B cells and ASCs. ASCs were the furthest along pseudotime, preceded by switched memory cells (Fig 2B). DN2 cells occupied the earliest pseudotime positions, consistent with their lower mutation burden in the BCR, indicating limited affinity maturation [38]. Non-switched memory and COMB cells spanned a wide range of pseudotimes, with non-switched memory skewing earlier than COMB cells. This suggests that COMB cells are further along the developmental trajectory than non-switched memory but less mature than switched memory cells (Fig 2B).

Monocle trajectory analysis further supported these findings (Fig 2C). Non-switched memory cells were positioned at the earliest pseudotime among memory subsets, while COMB cells appeared at a similar pseudotime to DN2 cells. DN2s were earlier along pseudotime than switched memory cells, with ASCs as the most mature population (Fig 2C). These analyses suggest that COMB cells are less mature than switched memory cells but represent a heterogeneous population spanning different levels of activation or differentiation. Additional studies are needed to clarify the relationships between these subsets and their precursors.

Given their distinct phenotype and increased abundance in RA (Fig 1E), we focused on COMB cells for further investigation. In addition to expressing *COCH, IGHE*, and *IGHG4* (Fig 2A), COMB cells exhibited high expression of activation-associated genes such as *SCIMP, TCF7, HOPX, TEX9, AIM2*, and *IL13RA1* (Fig 2D). Notably, *SCIMP* is involved in BCR signalling [39], while the other genes are linked to B cell activation, suggesting that COMB cells are activated.

To investigate whether COMB-like cells are present in diseased tissue, we created a gene module based on highly expressed COMB cell markers. Using scRNA-seq data from synovial RA samples previously generated in our lab [40], we found that the COMB module was enriched in synovial switched memory B cells (Fig 2E). These synovial switched memory cells also expressed high levels of *SCIMP, TCF7, HOPX, TEX9, AIM2, IGHE*, and *IL13RA1*, closely resembling circulating COMB cells. This suggests that COMB-like populations are present in RA synovium where they represent the main population of memory cells and may play a role in disease-specific immune responses.

This comprehensive analysis of circulating B cells reveals distinct developmental trajectories and activation states in RA compared to OA. Naïve B cell subsets dominate in OA, while RA is characterized by an enrichment of memory B cells, including Cochlin-Expressing Memory B (COMB) cells, an activated subset defined by unique marker expression (*COCH, SCIMP, TCF7, AIM2*) and increased abundance.

### COMB Cells as a Shared Immune Population Across Autoimmunity, Viral Infections, and Atopy

Having identified COMB cells in synovial tissue, we sought to determine shared gene expression patterns across different COMB cell populations and identify genes specifically upregulated in COMB cells from RA. We found that synovial COMB cells exhibited a gene expression profile more similar to blood COMB cells from RA than from OA. Among these, 33 genes were significantly upregulated across synovial RA COMB cells and blood RA and OA COMB cells, forming a core COMB gene signature (Fig. 3A).

**Fig 3.**
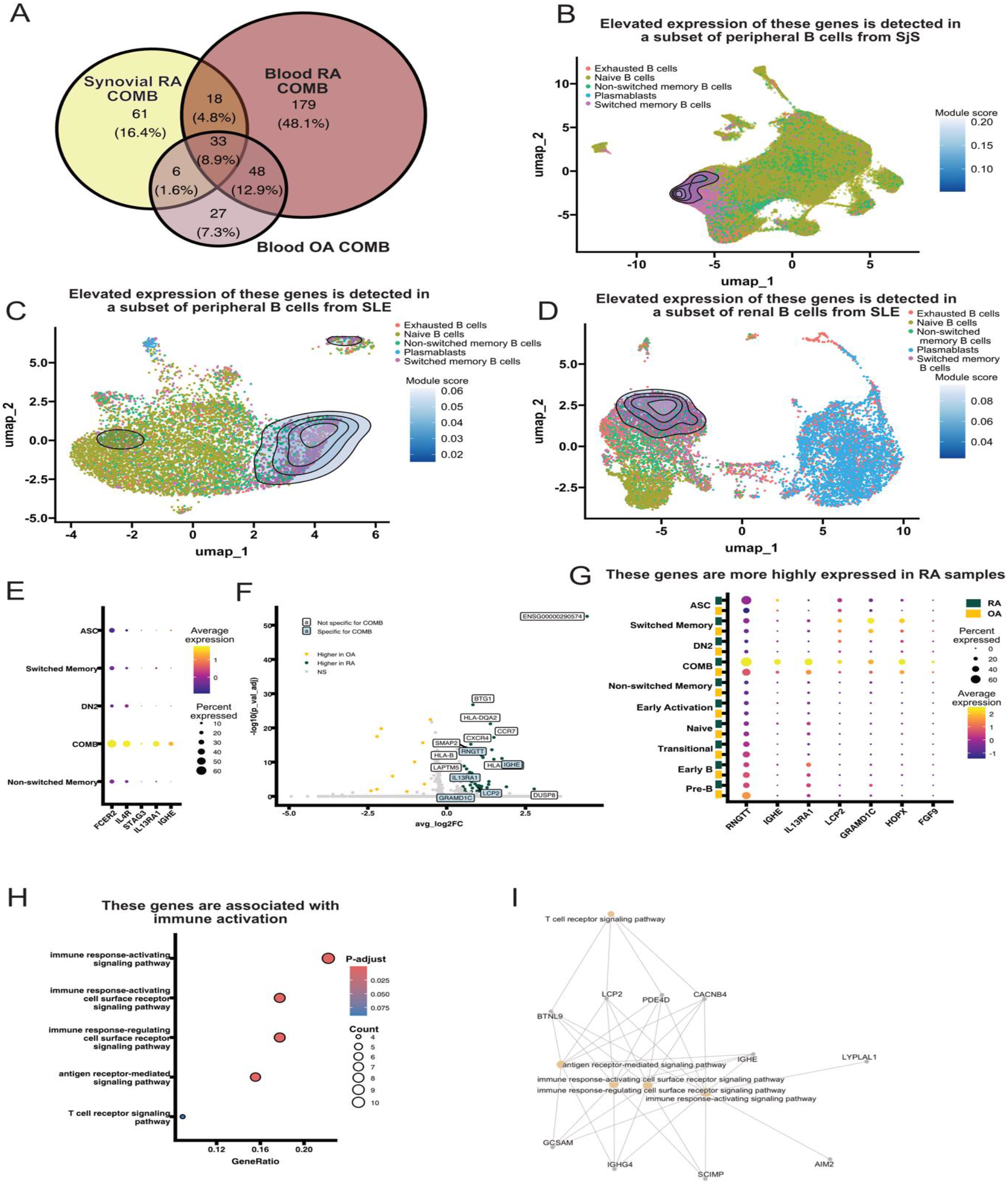
COMB Gene Expression Signatures in Autoimmune Diseases. **3A**. Venn diagram illustrating the overlap of significantly upregulated genes in COMB cells from different sources. Synovial RA COMB cells uniquely express 61 genes (16.4%), blood RA COMB cells uniquely express 179 genes (48.1%), and blood OA COMB cells uniquely express 27 genes (7.3%). Shared gene sets include 33 genes (8.9%) common to all three populations and 18 genes (4.8%) shared between synovial and blood RA COMB cells. **3B**. UMAP projection of scRNA-seq data from 236,088 peripheral B cells from healthy donors (n=4) and Sjögren’s syndrome (SjS) patients (n=26) with the module scores from the 33 COMB-specific genes, the contour overlay represents the position of the top 5% of module scores, identifying a subset of B cells with similar transcriptional profiles. **3C**. UMAP projection of scRNA-seq data from 8,141 peripheral B cells from healthy donors (n=6) and systemic lupus erythematosus (SLE) patients (n=9) with the module scores from the 33 COMB-specific genes, the contour overlay represents the position of the top 5% of module scores, suggesting the presence of B cells with COMB-like transcriptional profiles. **3D**. UMAP projection of scRNA-seq data from 10,968 renal B cells from systemic lupus erythematosus (SLE) patients (n=146) with the module scores from the 33 COMB-specific genes, the contour overlay represents the position of the top 5% of module scores, suggesting the presence of COMB-like cells. **3E**. Dot plot comparing the gene expression of COMB cells to other memory B cell subsets. COMB cells exhibit highly similar gene expression to CD45RB^lo^ cells, previously reported in viral infections, as well as IL4R^+ve^+FCER2^+ve^ memory B cells, which were described in atopic diseases. Key genes shared across these subsets include *IL13RA1, FCER2, IL4R*, and *IGHE*. **3F**. Volcano plot showing differential gene expression between COMB cells in OA and RA (yellow, log2FC is < −0.5 and adjusted p-value < 0.05; green, log2FC is > 0.5 and adjusted p-value < 0.05; grey, nonsignificant genes). Genes specifically upregulated in RA COMB cells, such as *IGHE, IL13RA1*, and *HOPX*, are highlighted, while differentially expressed genes shared across clusters are shown for comparison. **3G**. Dot plot illustrating the expression of key COMB-specific marker genes across B cell subsets in RA and OA samples. Dot size represents the percentage of cells expressing each gene, while colour indicates expression levels. **3H**. Gene Ontology (GO) term enrichment analysis of genes differentially expressed in COMB cells in RA. COMB cells in RA show higher expression of genes linked to immune response and immune signalling pathways compared to those in OA. Colour indicates adjusted p-value and size indicates the number of genes associated with each GO term. **3I**: Gene-concept network illustrating associations between the Gene Ontology (GO) term enrichment analysis and the relevant GO terms. Genes are represented as grey nodes, while GO terms are shown as beige nodes, highlighting connections to immune activation pathways.

Using this 33-gene signature, we investigated publicly available B cell datasets from other autoimmune diseases, focusing on primary SjS and SLE. In a SjS dataset, we identified a subset of peripheral B cells with high expression of COMB-specific genes [21] (Fig. 3B). Annotation using SingleR and the Monaco dataset revealed that these cells were predominantly within the switched memory B cell population, consistent with COMB cell identity. Similarly, we found a comparable cluster of COMB-like cells in peripheral B cells from SLE, also within the switched memory B cell subset cluster [22, 23] (Fig. 3C). The specificity of this COMB signature within switched memory B cells, along with its presence across multiple autoimmune diseases, strongly suggests that COMB cells play a broader role beyond RA.

To further explore whether COMB cells are present in diseased tissue, we examined scRNA-seq data from renal SLE patients using the ARK Portal (arkportal.synapse.org). We again observed high expression of COMB-specific genes within the switched memory B cell population (Fig. 3D), indicating that COMB cells are not only present in peripheral blood but also infiltrate affected tissues in SLE.

Next, we compared the gene expression of COMB cells to previously described switched memory B cell subsets where cochlin transcripts had been identified in single-cell analyses. First, we found that memory B cells with high *COCH* expression were identified in another autoimmune disease, neuromyelitis optica spectrum disorder [41]. Next, we identified a strong transcriptional similarity between COMB cells and CD45RBlo B cells, a population enriched in individuals with viral infections [42] (Fig. 3E). Additionally, another B cell subset, IL4R+FCER2+ memory B cells, described in atopic diseases [43], shared a highly similar transcriptional profile, expressing *HOPX, RNGTT, IL13RA1, FCER2, IL4R*, and *IGHE* (Fig. 3E). These findings suggest that COMB cells are not exclusive to the autoimmune diseases we investigated, but are likely involved in immune responses to viral infections and allergic conditions.

After establishing the presence of COMB cells across multiple conditions, we investigated how they differ between autoimmune (RA) and non-autoimmune (OA) environments. Differential gene expression analysis between COMB cells in RA and OA revealed distinct transcriptional differences (Fig. 3F). While some genes were differentially expressed across all cell populations, we focused on those specifically upregulated in COMB cells. Key marker genes, including *IGHE, IL13RA1*, and *HOPX*, were more highly expressed in RA than in OA (Fig. 3G), suggesting that COMB cells in RA exhibit a more activated state.

To explore this further, we performed Gene Ontology (GO) enrichment analysis on genes upregulated in RA COMB cells. This analysis showed that RA COMB cells express genes associated with immune activation, particularly immune response and immune signalling pathways (Fig. 3H). Many of the enriched GO terms overlapped, with several key genes— *BTNL9, LCP2, PDE4D, CACNB4, IGHE, SCIMP, IGHG4*, and *GCSAM*—contributing to these pathways (Fig. 3I). This suggests that COMB cells in RA are not only more activated but may also play a role in amplifying the immune response.

Overall, these findings demonstrate that COMB cells are a distinct population of switched memory B cells present in multiple autoimmune diseases, as well as in viral and allergic immune responses. In RA, COMB cells exhibit an activated transcriptional profile, suggesting that they may contribute to disease pathogenesis by enhancing immune signalling and activation.

### Single-cell ATAC-seq Analysis Reveals Distinct Regulatory Features of COMB Cells

After identifying COMB cells as a distinct B cell population, we investigated their regulatory mechanisms and potential drivers of proliferation. To do this, we performed single-cell ATAC-seq (scATAC-seq) to assess chromatin accessibility in different B cell subsets from OA and RA patients. We used the same patient samples as those for scRNA-seq and successfully sequenced two OA and three RA samples. Clustering analysis and projection of scRNA-seq data onto the scATAC-seq dataset revealed the same 10 major B cell clusters, confirming the presence of COMB cells in both OA and RA (Fig. 4A).

**Fig 4.**
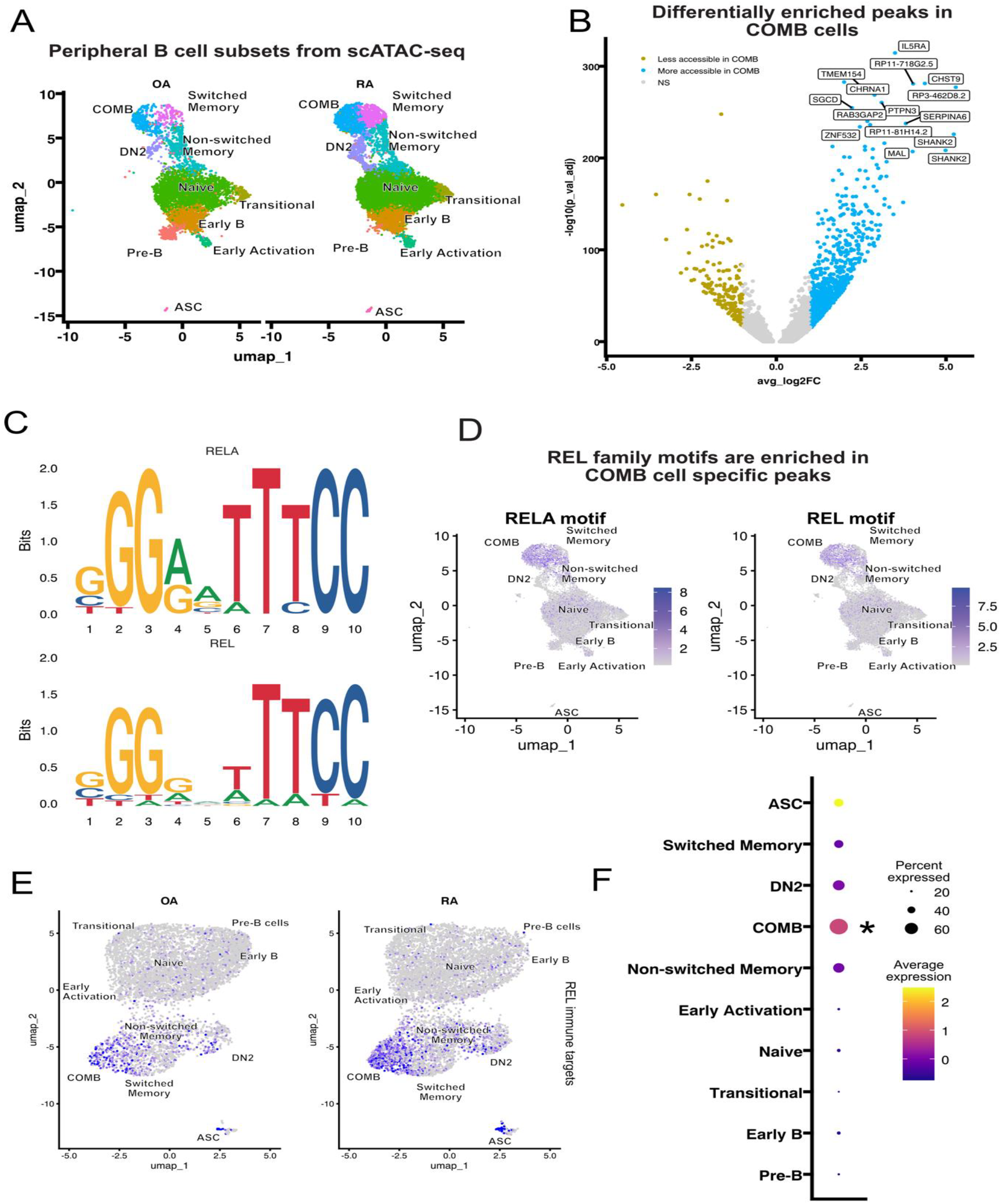
Single-cell ATAC-seq Analysis Reveals Distinct Regulatory Features of COMB cells. **4A**. UMAP visualization of peripheral B cell subsets identified by scATAC-seq showing distinct clustering of major populations with cells from OA samples on the left and RA on the right. **4B**. Volcano plot showing differentially accessible chromatin peaks in COMB cells. Accessible peaks are labelled by the closest gene. Genes such as *LISR4, SGCD*, and *SHANK2* exhibit increased accessibility in COMB cells compared to other subsets. **4C**. Sequence logos for RELA and REL consensus binding motifs enriched in chromatin regions specifically accessible in COMB cells. These motifs suggest involvement of NF-κB signalling pathways. **4D**. UMAP plots highlighting enrichment of RELA and REL motifs within chromatin regions accessible to COMB and switched memory B cell populations. **4E**. UMAP visualization showing the expression of REL-regulated immune genes across B cell subsets in OA (left) and RA (right) samples. A module of 13 immune genes regulated by c-Rel was identified, and their expression is significantly higher in the COMB cell cluster compared to other subsets. **4F**. Dot plot illustrating the expression of the REL-regulated immune gene module across B cell subsets. Excluding ASCs (that are distinct from COMB cells in their overall gene expression profile), COMB cells exhibit the highest expression of these genes, supporting the hypothesis that REL family transcription factors drive immune gene expression specifically in COMB cells. Dot size represents the percentage of cells expressing the module, while colour intensity reflects expression levels. * indicates a p-value < 0.05 following linear model testing, followed by Tukey test.

To identify differentially accessible chromatin regions in COMB cells, we compared their open chromatin profiles to all other B cell subsets. This analysis revealed that the *IL5RA* locus was particularly enriched in COMB cells, suggesting that these cells may be primed to respond to IL-5 (Fig. 4B). Notably, IL-5 levels are elevated in individuals at higher risk of developing RA, suggesting that this cytokine may play a role in COMB cell activation [44].

Next, we investigated the regulatory elements within the chromatin regions that were highly accessible in COMB cells. Motif enrichment analysis of these regions identified RELA and REL transcription factor motifs as significantly enriched in COMB-specific peaks (Fig. 4C). RELA and REL are genes encoding the NF-κB subunits RelA and c-Rel, respectively. These motifs belong to the NF-κB family, suggesting that NF-κB signalling is a key regulatory pathway in COMB cells. Given the sequence similarity between RELA and REL binding motifs, their concurrent enrichment further strengthens this finding (Fig. 4C).

To confirm the significance of these motifs at a single-cell level, we computed per-cell motif activity scores using chromVAR, which allows unbiased motif enrichment analysis across individual cells. This approach revealed that RELA and REL motifs were specifically enriched in COMB cells compared to all other B cell subsets, further supporting the involvement of NF-κB signalling in COMB cell regulation (Fig. 4D).

To understand the functional consequences of REL enrichment, we examined the expression of immune genes known to be regulated by c-Rel. We identified a set of 13 immune genes previously reported to be c-Rel targets, including *IGHG1, IGHG4, TNFA, IL12A, IL13, IL21, IL23A, CD40LG, TNFSF13B, CXCL10, LIG1, MIG*, and *FOXP3* [28]. Analysis of our scRNA-seq dataset confirmed that these genes were specifically expressed in COMB cells compared to other B cell subsets (Fig. 4E).

To further validate this, we created a module of these 13 immune genes and analysed their expression across different B cell populations. Excluding antibody-secreting cells (ASCs)— which strongly express *IGHG1* and *IGHG4*—we found that COMB cells exhibited the highest expression of the c-Rel-regulated immune gene module (Fig. 4F).

Together, these findings suggest that COMB cells are regulated by REL family transcription factors, with NF-κB signalling playing a crucial role in driving immune gene expression in COMB cells. This regulatory landscape further supports the idea that COMB cells contribute to immune activation and disease pathogenesis in RA.

## Discussion

Our study has identified a distinct subset of cochlin expressing memory B cells—COMB cells—that are consistently found in several autoimmune diseases, including RA, SLE, and Sjögren’s, with evidence of their presence in both the peripheral blood and inflamed tissues. These cells are transcriptionally and epigenetically distinct, likely regulated by NF-κB/REL family transcription factors, and functionally linked to immune activation and cytokine signalling.

To gain a deeper understanding of how B cells from RA patients are regulated we performed single-cell ATAC-seq (scATAC-seq) to assess chromatin accessibility and compared them to B cells from OA patients. This revealed that COMB cells exhibit distinct epigenetic profiles, with significant chromatin accessibility at RELA and REL binding motifs, suggesting that NF-κB signalling is a key regulatory pathway in these cells. Given that NF-κB activation is a hallmark of autoimmune disease, the enrichment of these motifs in COMB cells supports their role in perpetuating chronic inflammation.

Motif enrichment analysis identified RELA and c-Rel (REL) consensus motifs as significantly enriched in COMB-specific peaks, implicating canonical and non-canonical NF-κB pathways in their regulation. Additionally, we identified a set of NF-κB-regulated immune genes, including *IL12A, IL21, CD40LG*, and *TNFSF13B*, that are highly expressed in COMB cells. These findings suggest that NF-κB-dependent transcriptional programming contributes to the activation state and functional properties of COMB cells in autoimmunity.

A striking feature of COMB cells is their high expression of cochlin (*COCH*), a protein previously thought to be primarily extracellular in the cochlea [45] and follicular dendritic cells (FDCs) of lymphoid tissues [46]. Cochlin’s well-established role in lymphoid architecture and cytokine modulation suggests that its expression in B cells may have functional implications for immune activation and retention in inflamed tissues [47].

Recent findings on cleaved cochlin in innate immunity provide further insights into its potential role in adaptive immunity. Jung et al. (2019) demonstrated that cleaved cochlin (LCCL domain) enhances innate immune responses by sequestering bacteria, recruiting neutrophils/macrophages, and modulating inflammatory cytokine production (IL-1β and IL-6) [48]. These cytokines are key mediators of autoimmune inflammation in RA [49], SLE [50, 51], and SjS [52, 53], raising the question of whether cochlin-expressing B cells contribute to chronic inflammation via similar mechanisms.

Beyond cytokine modulation, cochlin undergoes enzymatic cleavage, generating LCCL domain fragments in response to inflammatory stimuli [47]. Given that RA synovial fibroblasts are major producers of aggrecanases [54], it is possible that COMB cell-derived cochlin is cleaved within inflamed tissues, generating LCCL fragments that amplify immune activation. This feedback loop could sustain inflammatory signalling, supporting B cell activation and contributing to disease chronicity.

Additionally, cochlin’s von Willebrand factor domains allow it to bind collagen-rich structures, particularly in cartilage and connective tissue [55]. We hypothesise that COMB cells localise to specific inflammatory sites—such as cartilage-adjacent pannus tissue in RA—where they may contribute to ongoing tissue destruction.

Our findings suggest that COMB cells’ pathogenic role in autoimmunity may be driven by both epigenetic regulation and cochlin-mediated immune amplification. The interplay between NF-κB transcriptional programming and cochlin-dependent cytokine modulation provides a novel mechanism through which B cells contribute to chronic inflammation and tissue damage.

Future studies should investigate whether cochlin cleavage in B cells generates immunostimulatory LCCL fragments, similar to its role in innate immunity, whether targeting cochlin or its cleavage products could offer therapeutic potential in autoimmune diseases, and whether NF-κB inhibition in COMB cells could modulate cochlin expression and reduce inflammatory cytokine production.

In summary, our study positions COMB cells as key players in autoimmune inflammation, linking NF-κB-driven transcriptional regulation, chromatin accessibility, and cochlin expression to their pathogenic function. The parallels between cleaved cochlin’s role in bacterial immunity and its potential function in autoimmunity suggest that cochlin-expressing B cells may contribute to immune dysregulation via cytokine amplification, immune cell recruitment, and tissue retention mechanisms. These insights provide a foundation for future therapeutic strategies targeting cochlin-mediated pathways in B cell-driven autoimmunity.

## References

1. Miller, F., The increasing prevalence of autoimmunity and autoimmune diseases: an urgent call to action for improved understanding, diagnosis, treatment, and prevention. Current opinion in immunology, 2023. 80.

2. Dörner, T., et al., B cells in autoimmunity. Arthritis Research & Therapy 2009 11:5, 2009. 11(5).

3. O’Dell, J.R., et al., Therapies for Active Rheumatoid Arthritis after Methotrexate Failure. New England Journal of Medicine, 2013. 369(4).

4. Abeles, I., et al., B Cell–Directed Therapy in Autoimmunity. Annual Review of Immunology, 2024. 42(Volume 42, 2024).

5. Jenks, S.A., et al., Distinct Effector B Cells Induced by Unregulated Toll-like Receptor 7 Contribute to Pathogenic Responses in Systemic Lupus Erythematosus. Immunity, 2018. 49(4).

6. Wing, E., et al., Frontiers | Double-negative-2 B cells are the major synovial plasma cell precursor in rheumatoid arthritis. Frontiers in Immunology, 2023. 14.

7. Weisel, F., U. Wellmann, and T.H. Winkler, Autoreactive B cells get activated in extrafollicular sites. European Journal of Immunology, 2007. 37(12).

8. Bagherzadeh-Fard, M., et al., The prevalence of thyroid dysfunction and autoimmune thyroid disease in patients with rheumatoid arthritis. BMC Rheumatology 2022 6:1, 2022. 6(1).

9. Ruacho, G., et al., Sjögren Syndrome in Systemic Lupus Erythematosus: A Subset Characterized by a Systemic Inflammatory State. The Journal of Rheumatology, 2020. 47(6).

10. Pasoto, S., V. Adriano de Oliveira Martins, and E. Bonfa, Sjögren’s syndrome and systemic lupus erythematosus: links and risks. Open access rheumatology : research and reviews, 2019. 11.

11. Shlomchik, M.J., et al., From T to B and back again: positive feedback in systemic autoimmune disease. Nature Reviews Immunology 2001 1:2, 2001. 1(2).

12. Calabrese, C.M., et al., Breakthrough SARS–CoV”2 Infections in Patients With Immune”Mediated Disease Undergoing B Cell–Depleting Therapy: A Retrospective Cohort Analysis. Arthritis & Rheumatology, 2022. 74(12).

13. Gray, M., et al., Apoptotic cells protect mice from autoimmune inflammation by the induction of regulatory B cells. Proceedings of the National Academy of Sciences of the United States of America, 2007. 104(35).

14. Miles, K., et al., A tolerogenic role for Toll-like receptor 9 is revealed by B-cell interaction with DNA complexes expressed on apoptotic cells. Proceedings of the National Academy of Sciences of the United States of America, 2011. 109(3).

15. Matsushita, T., et al., Regulatory B cells inhibit EAE initiation in mice while other B cells promote disease progression. The Journal of clinical investigation, 2008. 118(10).

16. Hao, Y., et al., Dictionary learning for integrative, multimodal and scalable single-cell analysis. Nature Biotechnology 2023 42:2, 2023. 42(2).

17. Aran, D., et al., Reference-based analysis of lung single-cell sequencing reveals a transitional profibrotic macrophage. Nature Immunology 2019 20:2, 2019. 20(2).

18. Angerer, P., et al., destiny: diffusion maps for large-scale single-cell data in R. Bioinformatics, 2016. 32(8).

19. Trapnell, C., et al., The dynamics and regulators of cell fate decisions are revealed by pseudotemporal ordering of single cells. Nature Biotechnology, 2014. 32(4).

20. Dann, E., et al., Differential abundance testing on single-cell data using k-nearest neighbor graphs. Nature Biotechnology 2021 40:2, 2021. 40(2).

21. Arvidsson, G., et al., Multimodal Single”Cell Sequencing of B Cells in Primary Sjögren’s Syndrome. Arthritis & Rheumatology, 2024. 76(2).

22. Mistry, P., et al., Transcriptomic, epigenetic, and functional analyses implicate neutrophil diversity in the pathogenesis of systemic lupus erythematosus. Proceedings of the National Academy of Sciences, 2019. 116(50).

23. Nehar-Belaid, D., et al., Mapping systemic lupus erythematosus heterogeneity at the single-cell level. Nature Immunology 2020 21:9, 2020. 21(9).

24. Xu, S., et al., Using clusterProfiler to characterize multiomics data. Nature Protocols, 2024. 19(11).

25. Stuart, T., et al., Single-cell chromatin state analysis with Signac. Nature Methods 2021 18:11, 2021. 18(11).

26. Wickham, H., ggplot2: Elegant Graphics for Data Analysis. 2016: Springer-Verlag New York.

27. Schep, A.N., et al., chromVAR: inferring transcription-factor-associated accessibility from single-cell epigenomic data. Nature Methods 2017 14:10, 2017. 14(10).

28. Gilmore, T.D. and S. Gerondakis, The c-Rel Transcription Factor in Development and Disease. Genes & Cancer, 2011. 2(7).

29. Bates, D., et al., Fitting Linear Mixed-Effects Models Using lme4. Journal of Statistical Software, 2015. 67(1).

30. Bretz, F., T. Hothorn, and P. Westfall, Multiple Comparisons Using R. 2016.

31. Campbell, K.S., et al., IgM antigen receptor complex contains phosphoprotein products of B29 and mb-1 genes. Proceedings of the National Academy of Sciences, 1991. 88(9).

32. LeBien, T.W. and T.F. Tedder, B lymphocytes: how they develop and function. Blood, 2008. 112(5).

33. Swaminathan, S., et al., BACH2 mediates negative selection and p53-dependent tumor suppression at the pre-B cell receptor checkpoint. Nature Medicine 2013 19:8, 2013. 19(8).

34. Swaminathan, S., C. Duy, and M. Müschen, BACH2–BCL6 balance regulates selection at the pre-B cell receptor checkpoint. Trends in Immunology, 2014. 35(3).

35. Palanichamy, A., et al., Novel Human Transitional B Cell Populations Revealed by B Cell Depletion Therapy. The Journal of Immunology, 2009. 182(10).

36. Sims, G.P., et al., Identification and characterization of circulating human transitional B cells. Blood, 2005. 105(11).

37. Sanz, I., et al., Challenges and Opportunities for Consistent Classification of Human B Cell and Plasma Cell Populations. Frontiers in Immunology, 2019. 10.

38. Cowan, G.J.M., et al., In Human Autoimmunity, a Substantial Component of the B Cell Repertoire Consists of Polyclonal, Barely Mutated IgG+ve B Cells. Frontiers in Immunology, 2020. 11.

39. Draber, P., et al., SCIMP, a Transmembrane Adaptor Protein Involved in Major Histocompatibility Complex Class II Signaling. Molecular and Cellular Biology, 2011. 31(22).

40. Wing, E., et al., Double-negative-2 B cells are the major synovial plasma cell precursor in rheumatoid arthritis. Frontiers in Immunology, 2023. 14.

41. Zhang, C., et al., B-Cell Compartmental Features and Molecular Basis for Therapy in Autoimmune Disease. Neurology® Neuroimmunology & Neuroinflammation, 2021. 8(6).

42. Priest, D.G., et al., Atypical and non-classical CD45RBlo memory B cells are the majority of circulating SARS-CoV-2 specific B cells following mRNA vaccination or COVID-19. Nature Communications 2024 15:1, 2024. 15(1).

43. Aranda, C.J., et al., IgG memory B cells expressing IL4R and FCER2 are associated with atopic diseases. Allergy, 2023. 78(3).

44. Chalan, P., et al., Analysis of serum immune markers in seropositive and seronegative rheumatoid arthritis and in high-risk seropositive arthralgia patients. Scientific Reports 2016 6:1, 2016. 6(1).

45. Bhattacharya, S.K., Focus on molecules: Cochlin. Experimental eye research, 2005. 82(3).

46. Py, B.F., et al., Cochlin produced by follicular dendritic cells promotes anti-bacterial innate immunity. Immunity, 2013. 38(5).

47. Rhyu, H.-J., et al., Cochlin-cleaved LCCL is a dual-armed regulator of the innate immune response in the cochlea during inflammation. BMB Reports, 2020. 53(9).

48. Jung, J., et al., Cleaved Cochlin Sequesters Pseudomonas aeruginosa and Activates Innate Immunity in the Inner Ear. Cell Host & Microbe, 2019. 25(4).

49. Kondo, N., et al., Cytokine Networks in the Pathogenesis of Rheumatoid Arthritis. International Journal of Molecular Sciences 2021, Vol. 22, Page 10922, 2021. 22(20).

50. Umare, V., et al., Effect of Proinflammatory Cytokines (IL”6, TNF”α, and IL”1β) on Clinical Manifestations in Indian SLE Patients. Mediators of Inflammation, 2014. 2014(1).

51. Linker-Israeli, M., et al., Association of IL-6 gene alleles with systemic lupus erythematosus (SLE) and with elevated IL-6 expression. Genes & Immunity 1999 1:1, 1999. 1(1).

52. Yang, H., Cytokine expression in patients with interstitial lung disease in primary Sjogren’s syndrome and its clinical significance. Am J Transl Res, 2021. 13(7): p. 8391–8396.

53. Huang, Y.-f., et al., The Immune Factors Involved in the Pathogenesis, Diagnosis, and Treatment of Sjogren′s Syndrome. Journal of Immunology Research, 2013. 2013(1).

54. Yamanishi, Y., et al., Expression and Regulation of Aggrecanase in Arthritis: The Role of TGF-β. The Journal of Immunology, 2002. 168(3).

55. Nagy, I., M. Trexler, and L. Patthy, The second von Willebrand type A domain of cochlin has high affinity for type I, type II and type IV collagens. FEBS Letters, 2008. 582(29).

