## Supplemental Table 1 and supplementary figure S1 for "Cochlin-Expressing Memory B (COMB) Cells Are Enriched in Autoimmune Diseases and Display a Distinct Activation Profile"

Supplementary Table 1. Antibodies used for B cell sorting with FACS

| Antibody (clone) | Fluorochrome | Isotype | Source | Dilution |
|---|---|---|---|---|
| Anti-human CD3 (HIT3a) | FITC | Mouse IgG2a, κ | BioLegend | 1:50 |
| Anti-human CD19 (HIB19) | PE | Mouse IgG1, κ | BioLegend | 1:50 |
| Viability dye | DAPI | | Sigma-Aldrich | 1:1000 |

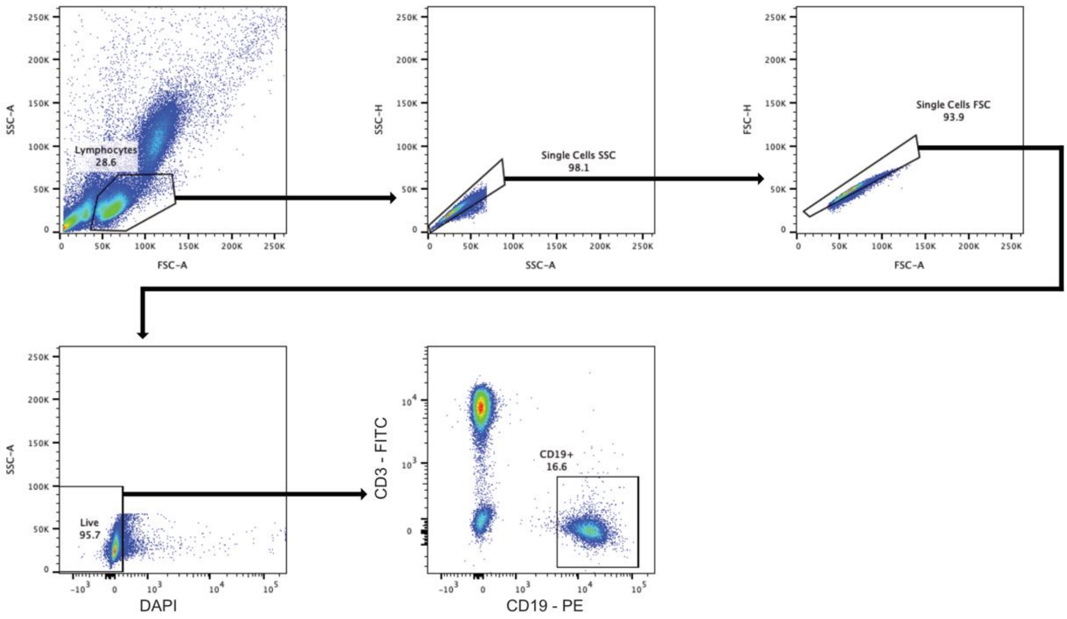


Supplementary Figure S1: Isolation and annotation of CD19+ve peripheral B cells.

A) Flow cytometry gating strategy for FACS sorting CD19+ve B cells from human PBMCs; representative plot from one samples.
